# Deep feature selection for Identification of Essential Proteins of Learning and Memory in Mouse Model of Down Syndrome

**DOI:** 10.1101/333849

**Authors:** Sara S. Abdeldayem, Mahmoud M. Elhefnawi

**Affiliations:** System and Biomedical Department, Faculty of Engineering, Cairo University, Egypt; The National Research Centre, Biomedical informatics and Chemoinformatics group, Cairo, Egypt

## Abstract

Down syndrome is a chromosomal abnormality related to intellectual disabilities that affects 0.1% of live births worldwide. It occurs when an individual has a full or partial extra copy of chromosome 21. This chromosome trisomy results in the overexpression of genes that is believed to be sufficient to interfere normal pathways and normal responses to stimulation, causing learning and memory deficiency. Therefore, by studying these proteins and the disturbance in pathways that are involved in learning and memory, we can consider drugs that would correct the observed perturbations, and therefore assist in enhancing the memory and learning. Here, from genes based on an earlier study that identified 77 proteins differentially expressed in normal and trisomic wild mice exposed to context fear conditioning (CFC), we provide a quantitative protein selection based on different feature selection techniques to select the most important proteins related to learning and memory. These techniques include Fisher score, Chi score, and correlation-based subset. In addition, a deep feature selection is utilized to extract high order proteins using deep neural networks. Three main experiments are carried out:studying the control mice’s response, studying the trisomy mice’s response, and studying the control-trisomy mice’s response. In each experiment, support vector machine classifier is used to assess these selected proteins ability to distinguish between learned and not-learned mice to the fear conditioning event. By applying the deep feature selection, fifteen proteins were selected in control mice, nine in trisomy mice, and seven in control-trisomy mice achieving distinguishing accuracies of 93%, 99%, 84% respectively compared to 74%, 78%, and 71% average accuracies of other selection methods. Some of these proteins have important biological function in learning such as CaNA, NUMb, and NOS.

## Introduction

Down syndrome (DS) is one of the most prominent causes of memory and learning deficiency [1]. There are around 3,000 to 5,000 children born annually with down syndrome [2]. DS is usually related to intellectual disability, and physical growth delays [3]. It is a chromosomal anomaly caused by the presence of a third copy, full or partial, of chromosome 21 (Hsa21). This can be due to an error in cell division, non-disjunction, in which a pair of 21st chromosomes fail to separate, resulting in an additional chromosome called trisomy 21 [2]. This Hsa21 encodes more than 500 gene models [4], where an unknown subset of these genes contributes in the learning deficit [5]. Overexpression of these genes may affect different biological processes and pathways, including brain development and function [6].

Along the past decades, the interest in the identification of DS learning deficits has been raised [7–13]. However, because of the lack of enough information about Hsa21 encoded genes, where functional information is available for less their half [5], it is logical to look for a disturbance in pathways that are critical to learning and memory, and then to consider drugs that would correct the observed perturbations. The benefit of studying pathway is that deep understanding of the functions of individual Hsa21 genes, their interactions, and contributions to brain function is not required [5]. The pathway disturbance can be gathered from experiments that each measures a small number of proteins. These experiments differ in some factors such as the performed task, the task protocol, the timing of the measurement of the proteins (i.e., after an hour and some minutes), or the region and method of the protein analysis [14]. For instance, Ahmed et al. [14] used the reverse phase protein arrays (RPPA) to analyze the levels of more than 80 proteins in mice exposed to Context Fear Conditioning (CFC) to provide a study of protein responses and interactions after normal learning. Higuera at el. [5] analyzed 77 proteins levels of control and trisomy mice (down syndrome) with and without learning simulations to study the effect of treatment, and to investigate the involved proteins in the learning process. They used self-organizing maps (SOM), an unsupervised clustering method using neural networks, the output clusters of classes are then processed to gather similar neighbors together. Then, the Wilcoxon test is used to investigate the significance of each protein in separating two classes. The significant proteins are then processed to get the final results using the intersection of the compared classes pairs. However, self-organizing maps can result in divided subclusters, and it does not provide a quantification of the protein selection results.

## Our Motivation and Contribution

Based on the work of [5] that used SOM to find the significant proteins in learning, our work is two-fold. First, on the contrary to SOM, we propose a quantitative approach to investigate protein expressions in order to identify biologically important differences in protein levels in mice exposed to CFC using machine learning feature selection algorithms. We examine about 77 proteins in subcellular fractionation in brain regions of wild type mice exposed to CFC to assess their relative learning, where part of these mice have been simulated to context shock with and without treatment using memantine, since it is currently used for treatment of moderate to severe Alzheimer’s disease (AD) and has been proposed for treatment of learning deficiency in DS [15, 16]. Second, we investigate different protein selection approaches to reflect the linearity nature of the selected proteins. We also utilize deep learning to mimic the nonlinearity of the underlying selection process.

The research is organized as follows: Section 2 discusses the dataset used as well as the methodologies we utilized in order to select the proper proteins. In section 3, we discuss the results of applying feature selection approaches on the protein expressions. Then we conclude in section 4.

## Materials and Methods

In order to detect the important proteins that contribute in memory and learning process, we use their expressions as features in different feature selection models. Fig 1 illustrates the workflow of the proposed approach. As the protein expressions act as features, we use four different feature selection models: Fisher score, Chi score, correlation-based approach, and Deep feature selection (D-DFS). Model validation and performance quantification is then performed on each model. The dataset and its acquisition, as well as each model, are discussed below.

**Fig 1.**
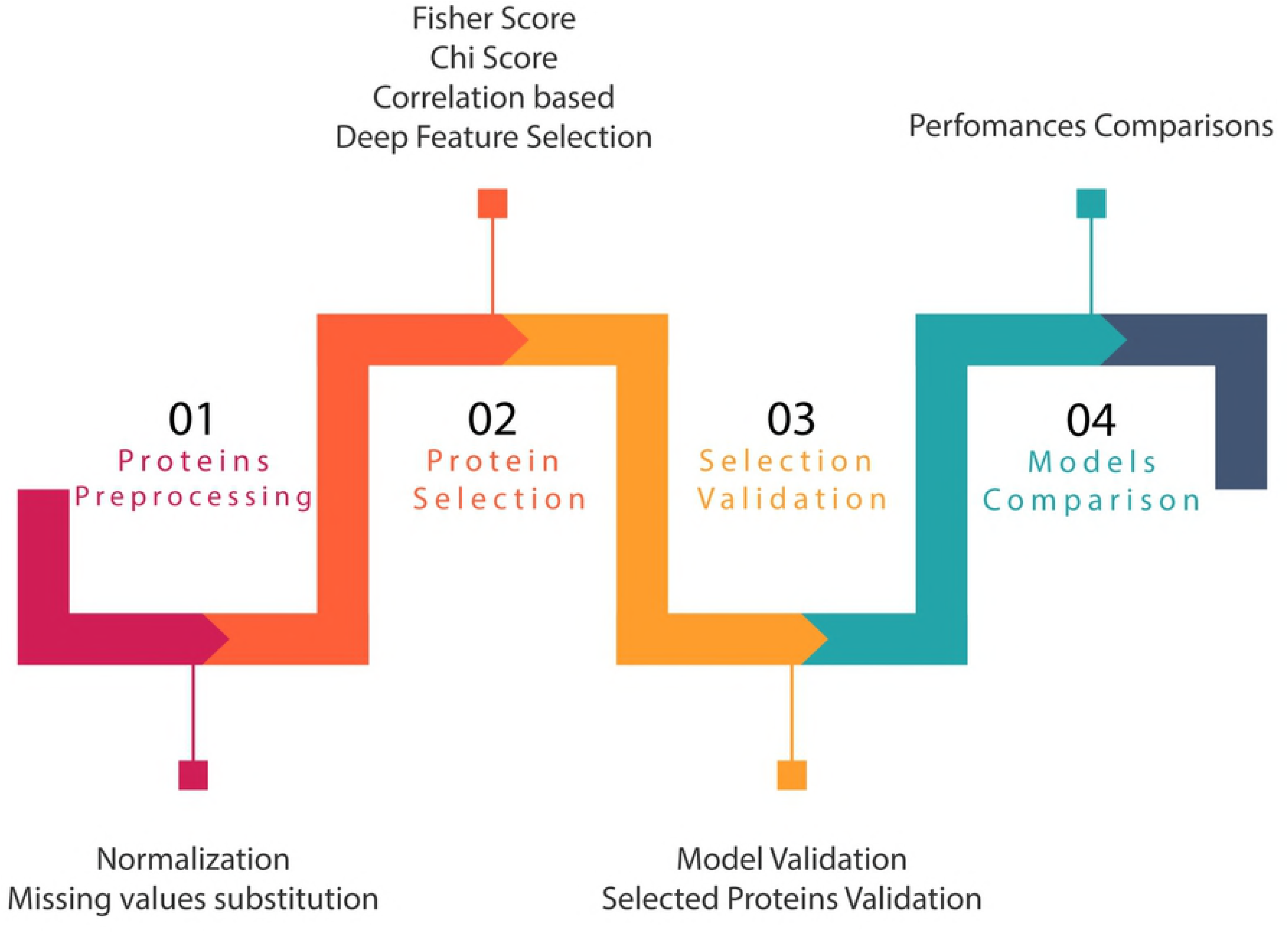
Workflow of the proposed approach.

## Abbreviation

Table 1 shows the abbreviation that we need to use along the paper. Though, these abbreviations will be defined in the text.

**Table 1.**
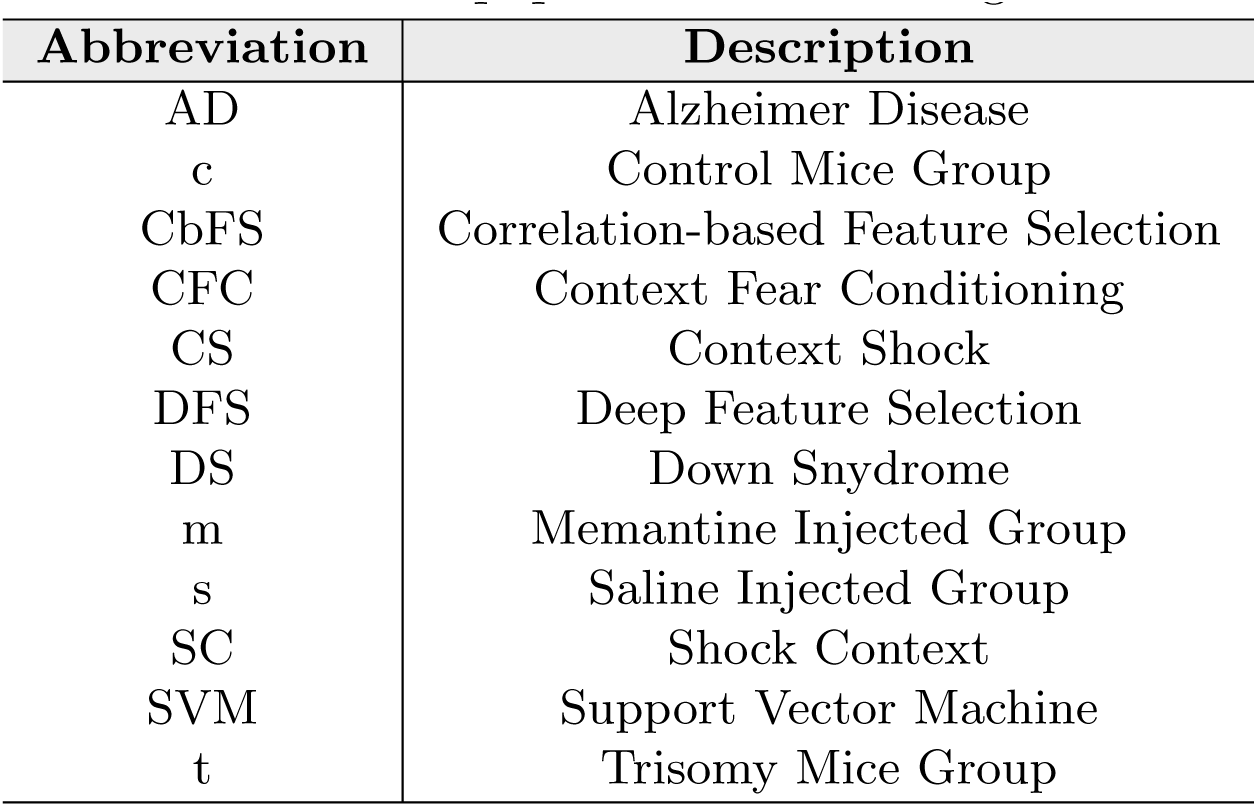
Abbreviations used in the paper and its meaning

## Dataset

In this study, we use a dataset of mice proteins expressions [14]. The dataset consists of the expression levels of 77 proteins/protein modifications that produced detectable signals in the nuclear fraction of the cortex. There are 38 control mice and 34 trisomic mice, for a total of 72 mice. In that experiment, 15 measurements were performed for each protein per mouse. Therefore, for control mice, there are 38×15, or 570 measurements, and for trisomic mice, there are 34×15, or 510 measurements. The dataset contains a total of 1080 measurements per protein, and each measurement can be considered as an independent sample [14]. Fig 2 shows the distribution of each mice group in the dataset.

**Fig 2.**
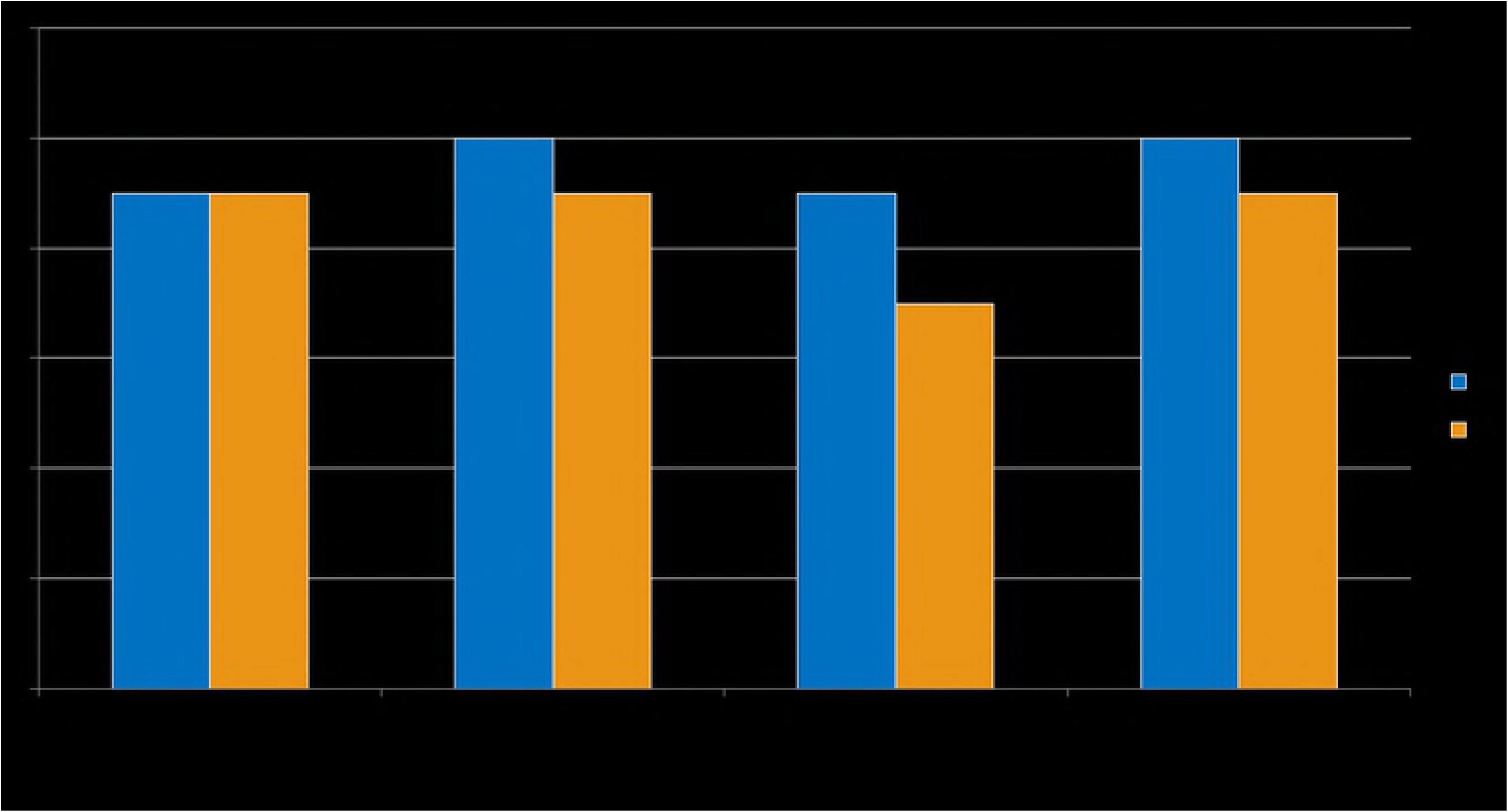
Mice dataset distribution.

### Dataset Acquisition

The dataset was a result of a context fear conditioning (CFC) experiment described in details in [14]. In that experiment, mice were first injected with memantine or the equivalent quantity of saline. Then, they were placed in a novel cage where they were allowed to explore for 3 minutes and then given an electric shock (2s, 0.7 mA, constant electric current). This group of mice had the chance to learn how to associate the context with the stimulus (electric shock) and accordingly is considered as context shock (CS) group. “Learning” here is meant by “freezing” after re-exposure to the context, where freezing is defined as a lack of movement except for respiration. Another group of mice, shock context (SC), were placed in the novel cage, and immediately given the electric shock, then they were allowed to explore for 3 minutes. Each group was studied under the effect of memantine and saline. Thus, a total of eight groups of mice were produced as illustrated in fig 3. A reverse phase protein arrays (RPPA) technique was then used to evaluate the levels of proteins and protein modifications in subcellular fractionation from hippocampus in the subject mice, where proteins that are relevant to CFC specifically, or to learning and memory and synaptic Alzheimer’s disease generally were chosen. However, as a reference, the name convention of the mice groups in this paper is as follow: (c/t)-(SC/CS)-(s/m), where c/t is for control or trisomy mice, SC/CS is for shock-context or context-shock mice, and s/m is either injection with saline or memantine. For instance, c-CS-s is for control mice that were exposed to context shock experiment while they were injected with saline, and t-SC-m is for trisomy mice that were injected with memantine and exposed to shock context experiment.

**Fig 3.**
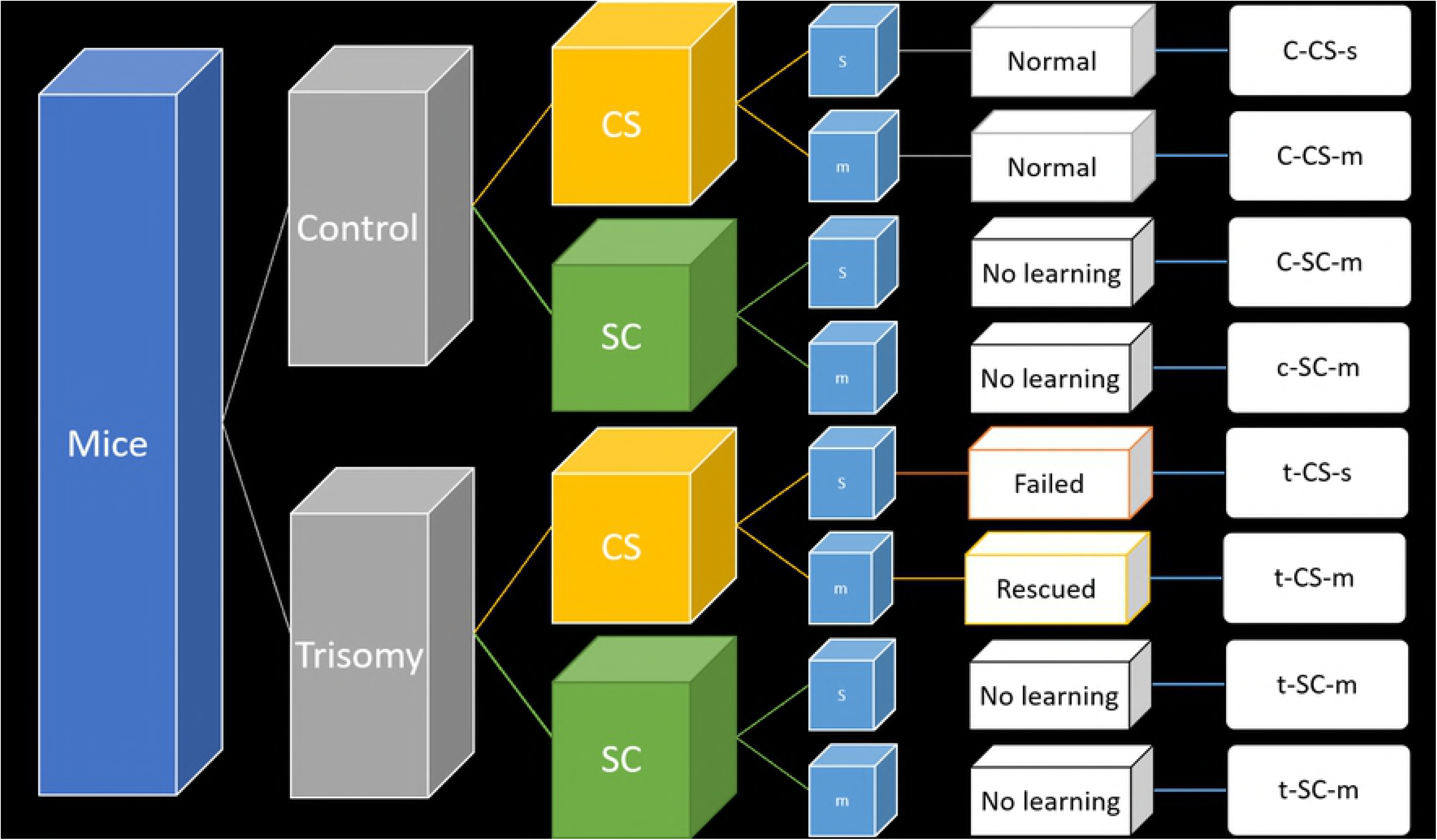
Mice proteins dataset description. At the first level, we have either control or trisomy mices. Each group is involved in context-shock and shock-context experiment, while injected by memantine or saline. The last level is the results of the mice’s responses.

### Data Processing

The proteins dataset used needed to be processed prior to investigation. First, the missing values problem in the dataset should be treated. Such that, for each missing protein value in any instance, we substitute it with the average protein expression in the instance’s corresponding class. Second, the dataset was normalized. Thus, the range of any proteins expressions became from 0 to 1 to avoid the domination of any protein of high expression values on other low expression proteins measurements.

### Protein Analysis

In order to investigate the proteins that relate to the memory and learning process, a feature selection algorithm is performed on the provided dataset, where the features are the protein expressions values. Features selection can be categorized into three main methods: wrapper methods, embedded methods, and, Filter methods.

Firstly, for wrapper methods [17], a search algorithm is applied in order to find the best subset of features. For instance, we can use the best-first search method [18], the random hill-climbing algorithm [19, 20], or the forward and backward passes [21] to add and remove features which can be considered as heuristic methods [22].

Secondly, for embedded methods, in which the accuracy of the system is measured, and efficient features are used during building the model. These methods are similar to the wrapper methods, but they are less computationally expensive, and prone to overfitting since the validation is done using cross-validation.

Thirdly, for the filter methods, a statistical measure is applied, and each feature is assigned a score. These scores are then ranked, and a threshold can be applied to eliminate the unnecessary features. In this paper, we will focus on this type of methods and consider both fisher, Correlation [23, 24], and Chi scores to investigate the importance of each protein in the learning process.

However, in some problems, we cannot simply apply these methods, since the data is more complex, and the system is nonlinear. Thus, we need to learn higher-level features in a way that can deal with nonlinearity. Deep neural networks have been used in these cases to improve the performance significantly, where nonlinearity and complexity of the system can be handled. In this paper, we employ Yifeng et al. [25] proposed method of feature selection using deep neural network modification.

In order to investigate the proteins that are important in the memory and learning, we divide the data into two main classes: mice that learned and mice that did not learn as in fig 4. First, we study the control mice group to know which proteins are responsible for the learning in case of control mice. For instance, in the first experiment we have two classes: c-CS-s and c-CS-m versus c-SC-s and c-SC-m. This will allow us to study the effect of the context shock or normal learning. Second, trisomy mice, we divide the trisomy classes into two classes, mice who rescued due to context shock and memantine, and those who failed to learn or did not learn due to the absence of a stimuli. Thus, we have t-CS-m versus t-CS-s, t-SC-s, and t-SC-m, where we can explore the effect of the rescued learning or the effect of context shock with memantine. Third, both the control and trisomy mice types are considered in the feature selection problem having two classes: c-CS-m, c-CS-s, and t-CS-m versus c-SC-s, c-SC-m, t-CS-s, t-SC-s, and t-SC-m. This will allow us to explore the difference between the normal learning in the control mice and memantine injected mice, and the not-learned mice due to lack of either context shock or memantine.

**Fig 4.**
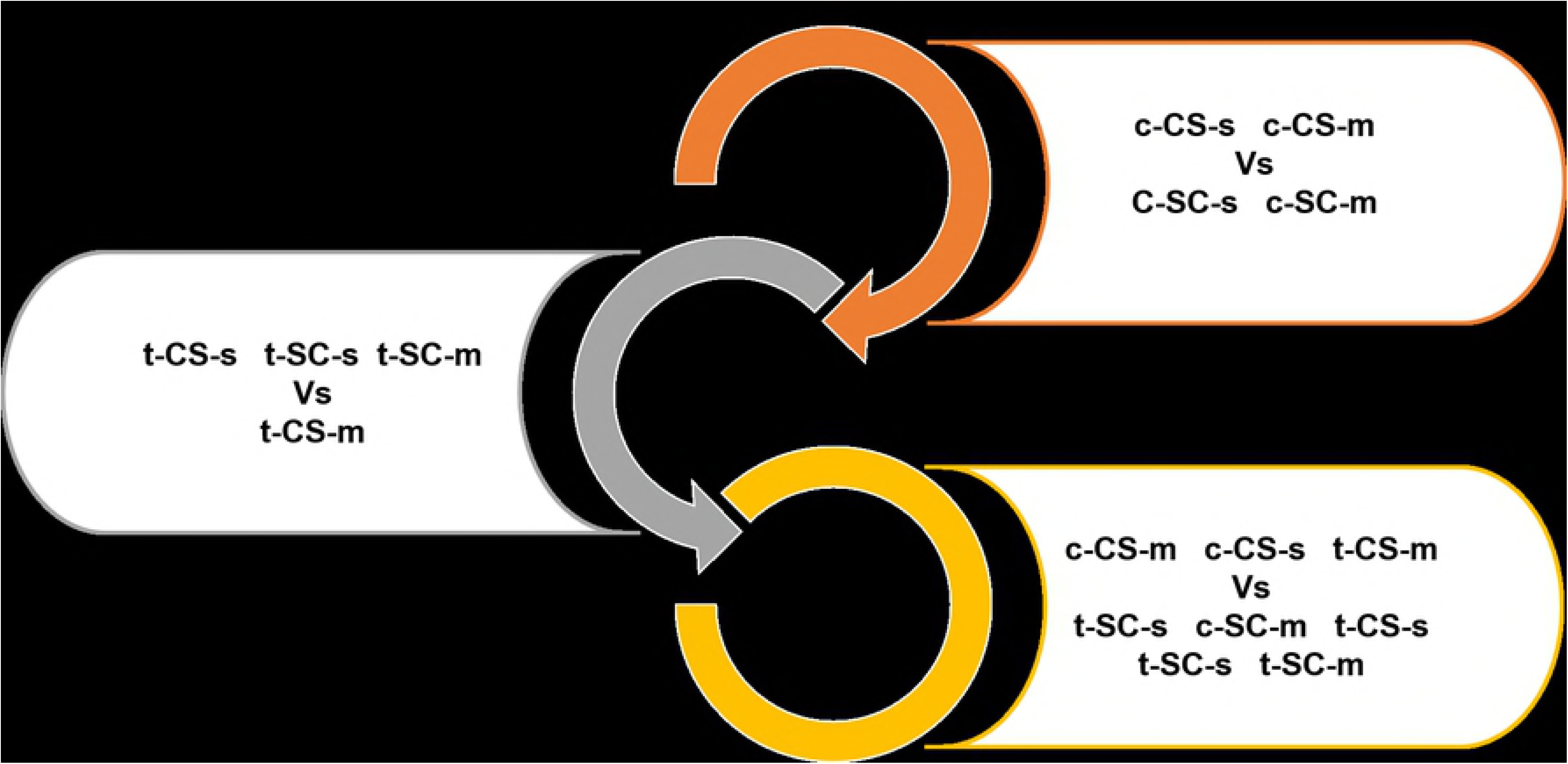
Proposed experiments of mice protein selection and classes involved.

## Fisher Score

Fisher score [26] is one of the most common supervised feature selection methods. It assigns a score to each feature independently according to the Fisher criterion. The features’ scores are then ranked, and the highest scores are selected. These scores depend on two measures; the within-class and between-classes distances. The within-class measure *S*_*w*_ expresses the distance between instances for the same class, while *S*_*b*_ represents the distance between the different classes as given by equations (1) and (2).

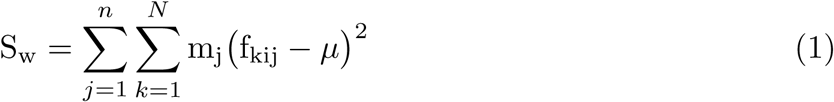

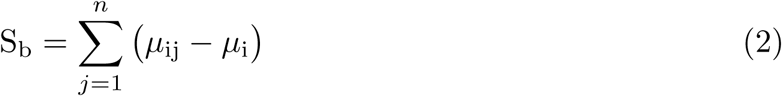

where *N* is the number of instances or observations, *j* is the class index up to n classes, *m*_*j*_ is the number of features in class *j*, *μ*_*i*_ is the mean of feature *i* along all classes, and *f*_*kij*_ is the value of feature *i* for observation *k* in class *j*.

The F score is then the ratio of *S*_*w*_ to *S*_*b*_ as in equation (3). This means that the higher F score, the better the feature will be.

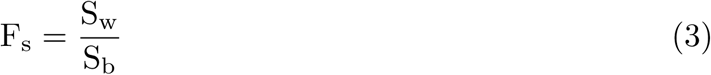

## Chi Score

Also called *χ*^2^ test or “goodness of fit” statistic. It reveals how likely it is that an observed distribution is due to chance, and quantify how well the observed data distribution fits compared to the expected distribution if the variables are independent [27]. Thus, the higher the Chi score, the better the feature classifies the data.

### Correlation-based Feature Selection (CbFS)

This method of feature selection’s central hypothesis is that good feature sets contain features that are highly correlated with the class, yet uncorrelated with each other [28]. It uses heuristic search methods to find the best features subset as in equation (4).

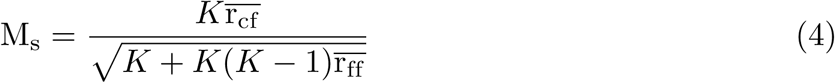

where *M*_*s*_ is the heuristic or merit of feature subset *S* that contains *K* features, r_cf_ is the mean feature-class correlation, and r_ff_ is the average feature-feature inter-correlation. The algorithm first computes the correlation factor of each feature. Then for every combination, the algorithm calculates its merit, and search for the best subset.

### Deep Feature Selection (DFS)

Deep neural networks are models structured by multiple hidden layers with nonlinear activation functions. It overcomes some problems of sparse linear models as it can handle nonlinear data, deal with complex data, and analyze data of higher orders according to the number of hidden layers. Since the weights of the nodes represent the importance of input values or the proteins, it can be used for sparsing of the features, where non-zero weights only can be considered. Usually, the sparsing and finding the weights of the network is done using a backpropagation method to find a solution of the following objective function:

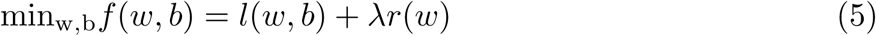

where *w* are the weights, *b* are the nodes’ biases, *l*(*w, b*) is the loss function (could be log likelihood, logistic logs, or squared loss), *r*(*w*) is the regularization term, and *λ* is the regularization factor that weights the regularization versus error.

Feature selection using neural network can be done in several ways. For instance, an elastic net can be used where the input is connected to one hidden layer and then the output layer (output nodes are equal to the number of classes) [25]. However, as we used only one hidden layer, the level of complexity concerned in this case is not high.

In shallow feature selection S-DFS, only one hidden layer exists in the network, and one layer is added in the input level that has the same number of nodes as the input. This additional layer’s weights are our concern, as only non-zero or above threshold weights are taken in the features selection as illustrated in Fig 5 (a). The output layer of the network is a softmax layer having a number of nodes equal to the number of classes. On the other hand, deep DFS, D-DFS, uses multi hidden layers based on the concept of Multilayer perceptron (MLP) [29]. This model is very useful in the systems of high complexity where data separation cannot be done linearly. In the same manner, an additional layer is added after the input layer that has the same number of nodes as the inputs (in our case proteins) as shown in Fig 5 (b). This will allow us to learn deeply about the proteins and their relations to each other. However, we will call the D-DFS as DFS for convenience.

**Fig 5.**
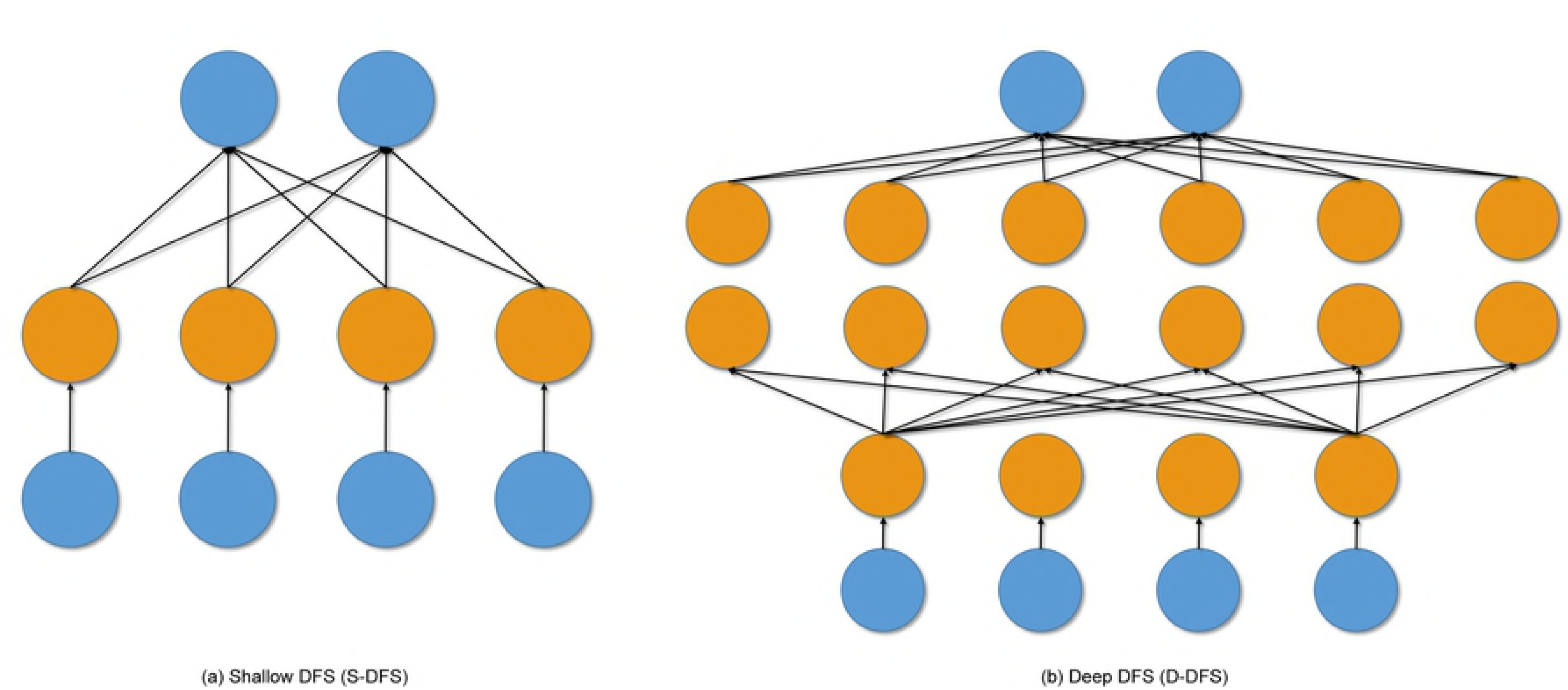
Neural Network configurations. (a) Shallow DFS (S-DFS). (b) Deep DFS (D-DFS).

In the DFS system, two regularization terms are used; one for the additional added layer and the other is for the hidden network layer. Thus, the objective function in equation (5) proposed by [25] will be like:

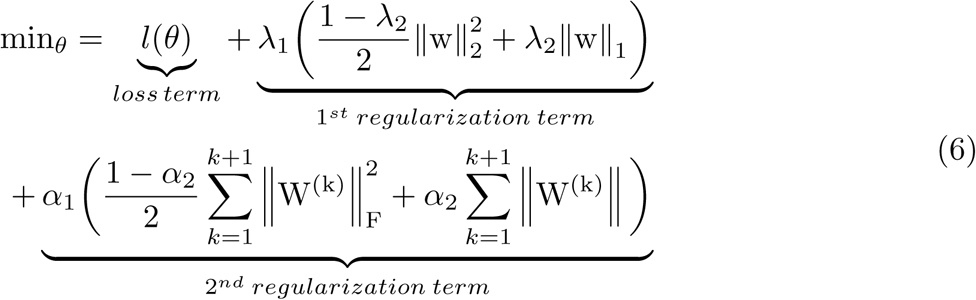

The equation can be explained as follows:

1. The loss term in our case is the log-likelihood, and it is given by:

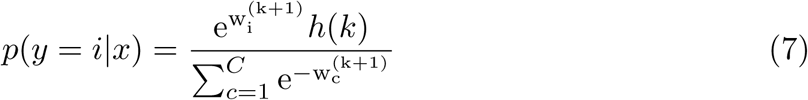

where *y* is the output, *x* is the input, *w*(*k* + 1) is the weight of the *k*^*th*^ layer, *i* is node index. Since the last layer is a softMax layer, its loss function will be like:

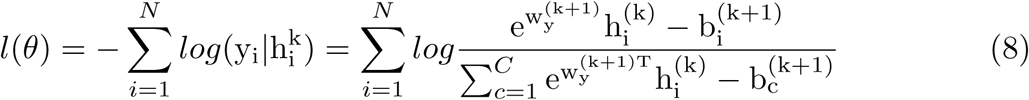

2. The first regularization term controls the tradeoff between sparsity of the weights and the smoothness of the network, where *λ*_2_ is a user-specified parameter, and can take range from 0 to 1.

3. The second regularization term in the equation is used to reduce the complexity of the model, and sparsing for the hidden net. Another effect of this term is to avoid the shrinking of weights in the input weights layer (the layer after the input layer) resulting having wi weights very small, and W weights large.

In order to find the solution to equation (6) we can utilized the gradient descent [30]. It is a first-order iterative optimization algorithm, where in each iteration it takes steps proportional to the negative of the gradient of the function at the current iteration.

### Selection Validation

For the purpose of quantifying and assessing the chosen proteins, we use classification models to measure the ability of these proteins to distinguish between the two classes studied. For instance, by selecting group X of proteins, the performance measures of the classifiers are reported. Thus, the higher performance, the better the protein group. We use three common measures named accuracy, sensitivity, and specificity that are defined as follow:

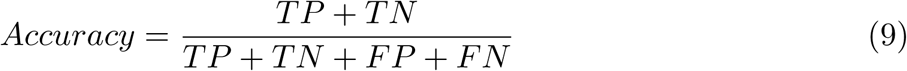

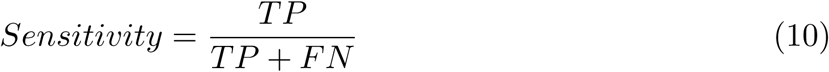

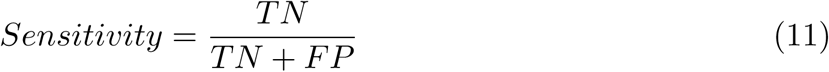

where TP is the true positive or the portion of positives that are correctly identified, TN is the true negative or potion of negatives that are correctly identified, FN is the false negative or the portion of positives that are identified as negative, and FP is the false positive or the portion of negatives that are classified as positive.

In order to validate the model, we use cross-validation technique. Using this validation technique, we can assess the generalization ability of the classification model. This generalization can be seen as the ability to make new predictions for data that has not already seen. In k-fold cross-validation, the original data is randomly partitioned into k equal size parts. Then for each iteration of the K iterations, a single subsample is used as a testing data for the model, and the remaining k-1 subsamples are used as training data. This cross-validation is utilized to avoid any over-fitting that might be encountered during the classification.

The classifier that we used in this paper is linear support vector machine (SVM) [31, 32]. The main idea of SVM is to find the optimal separating line between the data points or features to distinguish the classes. For instance, in a two-class SVM, the classifier searches for the closest feature points, called support vectors, where as the perpendicular to the line connecting these support vectors can be considered as the separating line between these two classes.

## Results and Discussion

Using the proteins dataset, the proposed protein selection algorithms were done using SVM in order to assess the performance of separation of each protein set. A five-fold cross-validation was utilized in all the experiments. First, the control mice data were divided into two classes: c-CS-m and c-CS-s, versus c-SC-m and c-SC-m. Second, the trisomy mice were divided into two classes dividing the t-CS-m versus the rest of trisomy classes. Finally, the proteins that affect the whole dataset discrimination were analyzed. In the DFS, a data separation was used having 70% as training data, 15% for validation data, and 15% for testing data.

## Control Mice Proteins Analysis

The fisher scores of the 77 proteins are illustrated in appendix 1. The SOD1 protein, superoxide dismutase 1, scored the highest F-score of 1.63 meaning that this protein has a high correlation with the learning and memory processes, which is supported by the results of [33]. The pPKCAB protein comes in the second place having a score of 1.28, followed by calcineurin CaNA that proved to influence both learning and reversal learning [34]. On the other hand, CAMKII, RRP1, GluR4, pRSK, CREB, GluR3, and RSK scored the lowest scores reflecting that these proteins do not affect the learning process in control mice according to the Fisher criterion.

The influence of the proteins sorted according to the Fisher score using SVM classifier is illustrated in Fig 6 (a), where the initial number of proteins is 1 and a step of 1 protein added was used to analyze the effect of the proteins. The maximum accuracy that can be achieved is 92.7% using all the proteins. However, the first 30 proteins span for 90% accuracy, having 47 proteins that only adds about 2% accuracy.

**Fig 6.**
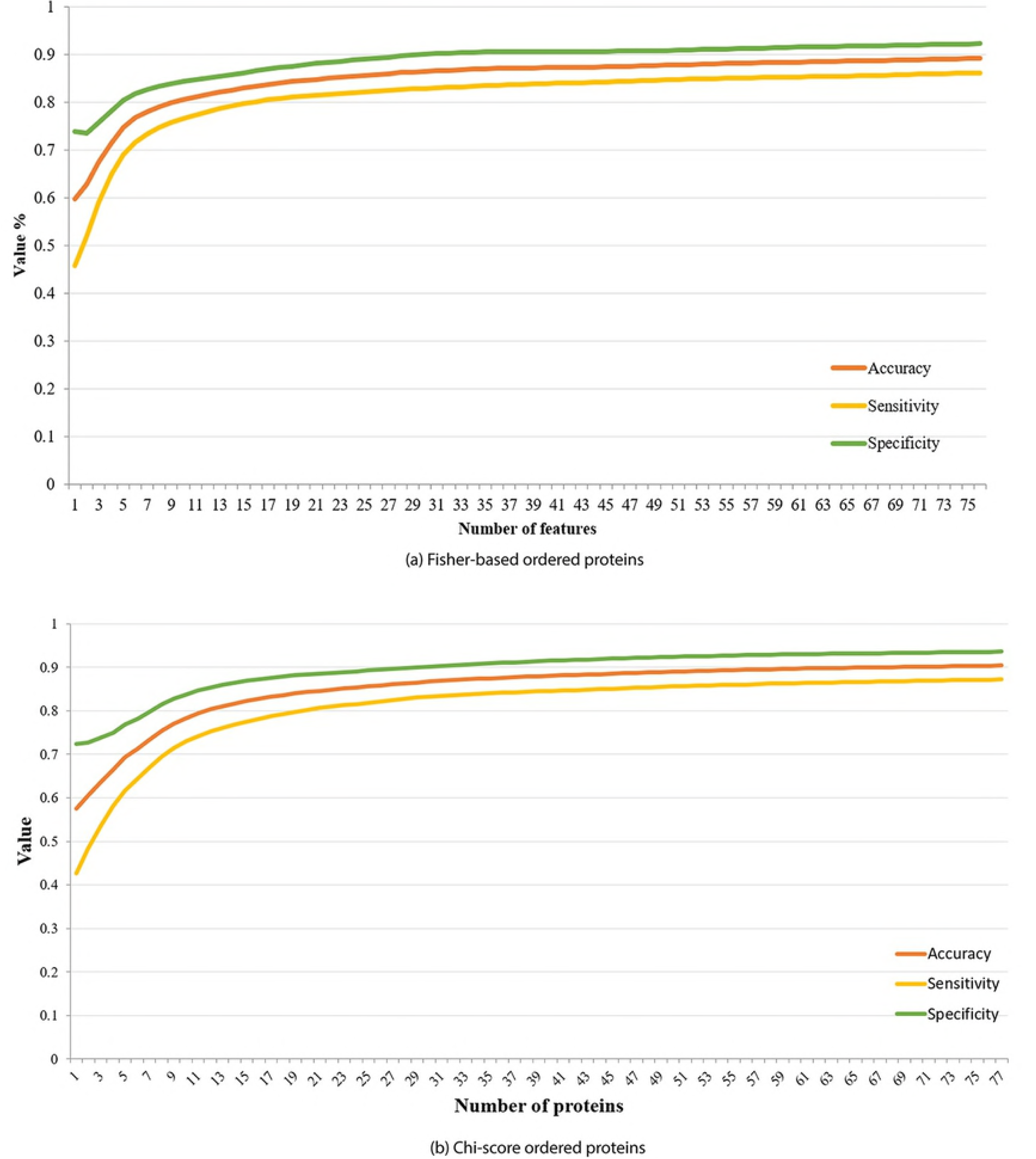
SVM Classification system performance in Control mice using different number of features sorted by (a) Fisher, (b) Chi scores.

The Chi score was then tested for the control mice classes as shown in appendix 1. It can be seen that, again, the SOD1 protein has the highest score of 695.5 followed by pPKCG with 614.3 score. We can also notice that CREB, pNR2B, and GluR3 have scores near zero, where they also gained the lowest scores in Fisher test. Hence, the Chi test is consistent with the results from the Fisher criterion. The proteins are then ordered, and the SVM classification was done on the data having a maximum accuracy of 91%, 78% sensitivity, and 93% specificity as shown in Fig 6 (b). As illustrated, the accuracy starts to settle after involving 33 proteins in the classification achieving 88% accuracy.

In order to test the performance of correlation-based feature selection analysis, we used the greedy stepwise algorithm for the search process starting with no attributes. The best merit found was 0.648. This approach presented a best subset of 17 proteins (no order): pCAMKII, pERK, PKCAB, AKT, SOD1, P38, pNUMB, pGSK3B, pPKCG, AcetylH3K9, ARC, nNOS, Ubiquitin, SHH, pCFOS N, H3MeK4, CaNA. Applying classification using only these 17 proteins yields an accuracy of 90%, sensitivity of 78%, and specificity of 93%. The SOD1, CaNA, and Ubiquitin scored the highest scores in both Fisher and Chi and was involved in the correlation best subset. On the other hand, pPKCG, pERK, H3Mek4, and CANA obtained a high score on both tests but were not selected by the correlation test.

In the deep feature selection, we used two hidden layers with a network configuration of 77, 100, 64, 2 nodes. The validation set was used to select the network parameters having *λ*_1_ = 0.01, *λ*_2_ = 1.0, learning rate *η* = 0.1, *α*_1_ = 0.0001, *α*_2_ = 0.6. The 77 weights of the added layer are shown in appendix 1. It can be shown that only 25 proteins have non-zero values. However, in order to get the efficient selected proteins, we applied thresholding equal to 5% of the maximum weight. This resulted in 15 proteins only: CREB, ERK, JNK N, PKCAB, CAMKII, Bcatenin, SOD1, pMTOR, DSCR1, pS6, BAX, NOS, Ubiquitin, H3MeK4, CaNA. Although some of the selected proteins may not have high scores in the previous selection techniques, these proteins have a large influence on higher orders because of the deep neural network. Though, SOD1, CaNA, and Ubiquitin have high values in all the techniques employed in control mice.

To assess the performance of the DFS, Fig 7 shows the comparison of the DFS with Fisher and Chi performance accuracies. We used a range of *λ*_1_ from 0.01 to 0.3 with a step of 0.001 to evaluate the number of selected proteins versus the accuracy. The DFS outperformed the SVM classification at the same number of proteins. In addition, DFS has steeply reached a good accuracy using less number of proteins. For example, having twelve proteins using the DFS results in an accuracy of 93% while the Fisher and Chi have 76% and 74%. This might be an indication of nonlinearity in the system and the need to investigate higher-level features such as in DFS. The DFS resulted in a cluster of low number of proteins because of the sparsity of the second layer.

**Fig 7.**
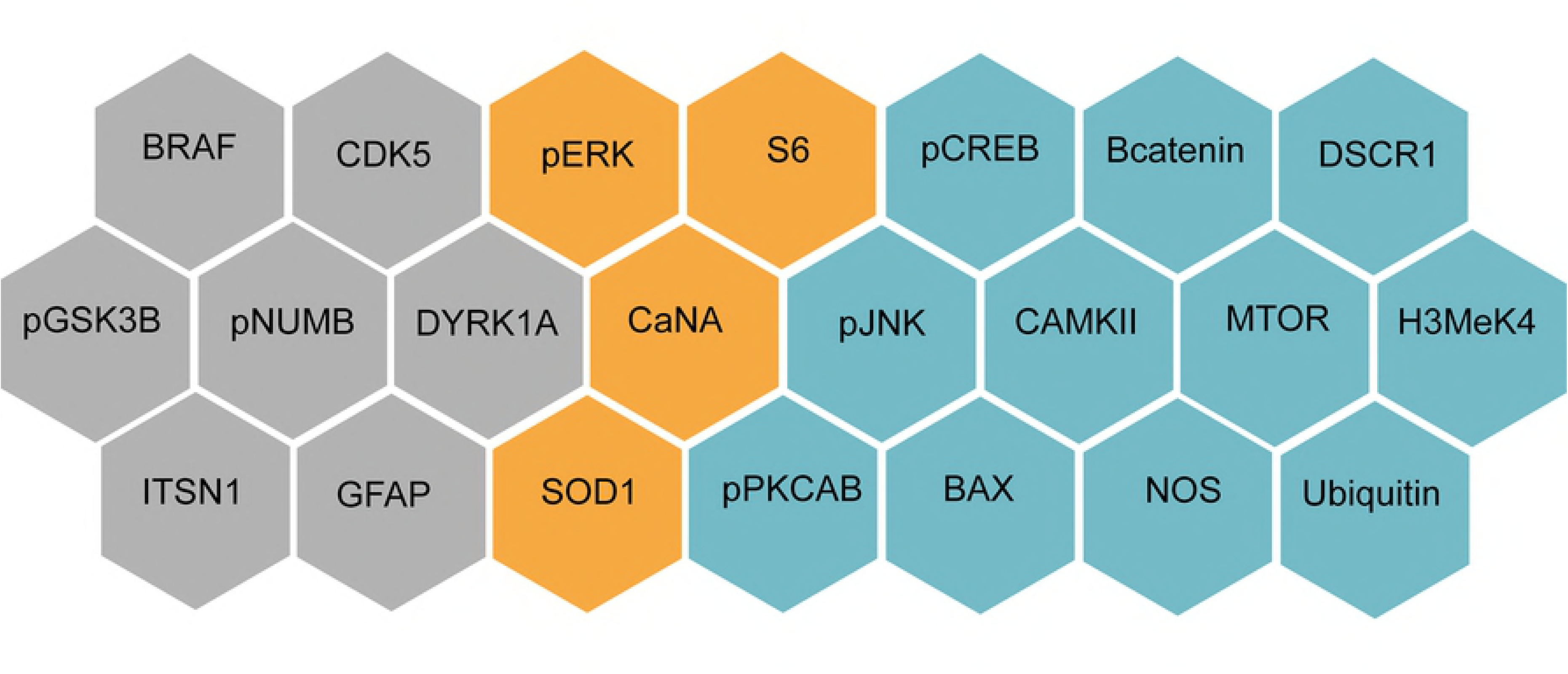
Performance of classification using different selection techniques in control mice using different number of proteins.

Comparing the proteins selected by the DFS and SOM [5], fig 8 shows the proteins selected by both algorithms illustrating the common proteins. Among the selected proteins pERK, S6, CaNA, and SOD1 were common. In order to assess our proteins, linear SVM was used with five cross-validation. The distinguishing accuracies of the five iterations in illustrated in Fig 9 using both sets of proteins selected by our approach and [5]. The proteins selected in our study show less variance in the accuracy with maximum and average accuracies of 91% and 87.19% respectively compared to 85.08%

**Fig 8.**
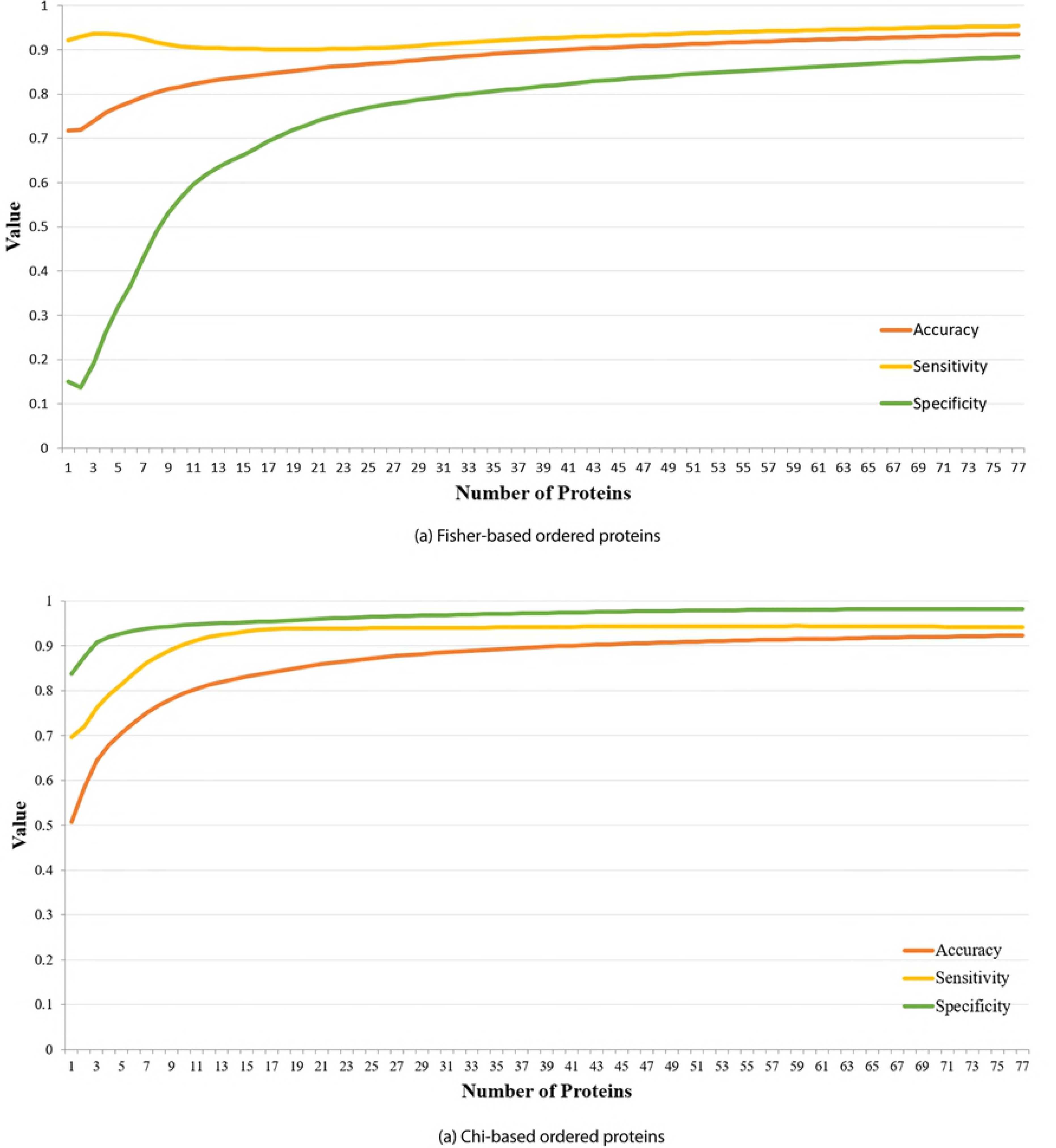
Proteins related to memory and learning from [5] and our approach in control mice group. Gray: proteins selected by [5]. Blue: proteins selected by DFS. Orange: common proteins. and 78.25% from [5]. Therefore, our selected proteins perform a better distinction between the two mice groups that are with and without learning experience.

**Fig 9.**
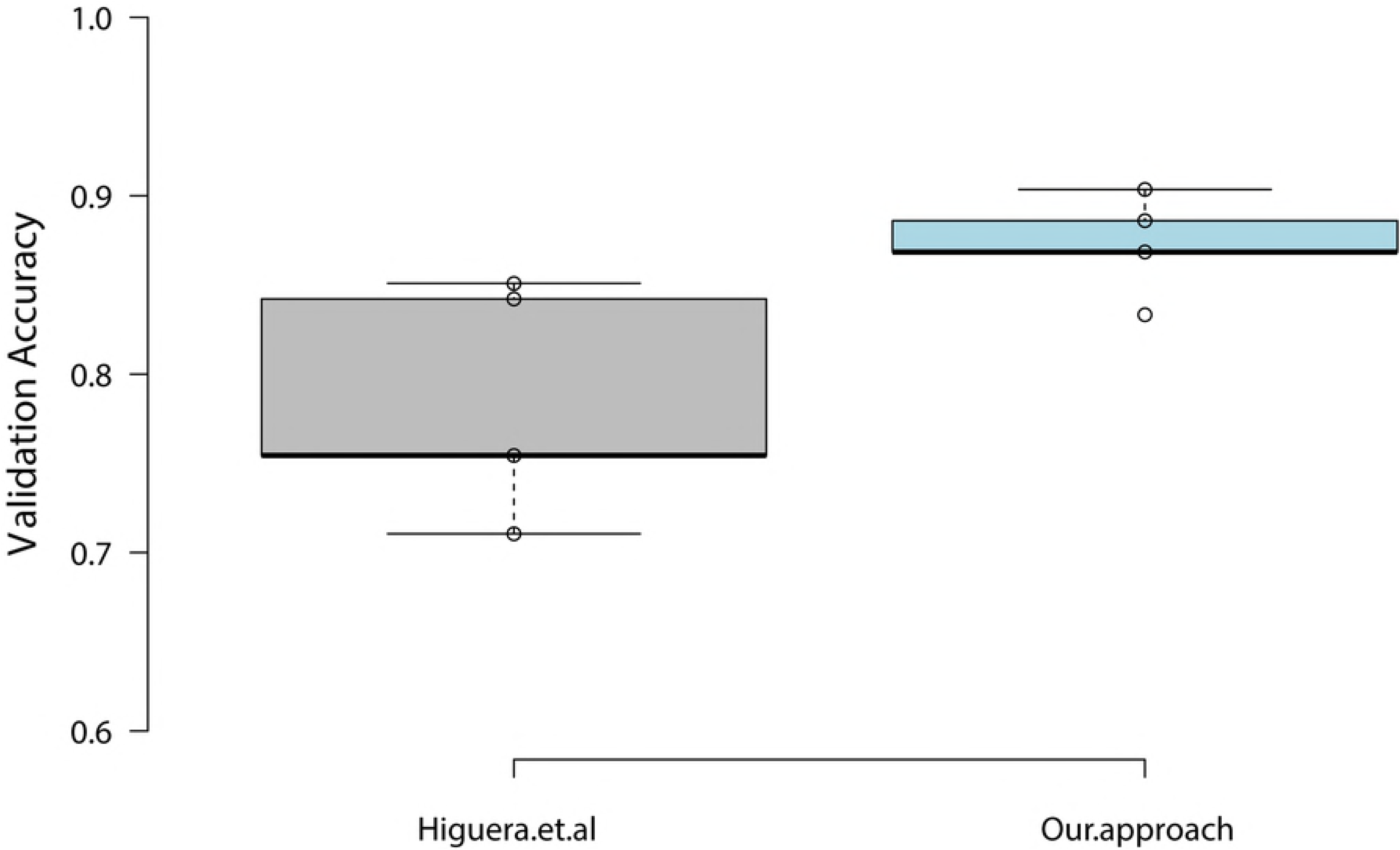
Boxplot of the SVM five cross-validation using the selected proteins in DFS and SOM in Control mice.

## Trisomy Mice Protein Analysis

The same algorithms were performed on the trisomy mice. Data was separated into two groups; a group which represents learning such as t-CS-m, and groups that failed to learn like t-CS-s, t-SC-m, and t-SC-s. The Fisher scores of the proteins can be shown in appendix 2 followed by an SVM classification performance assessment in Fig 10 (a). In contrast to the control mice, SOD1 was the fifth protein, while ARC and pS6 had the highest scores. The ribosomal protein pS6 has an impact on the short memory formation, storage, and retrieval as presented in Giese at el. [35]. On the other hand, we have a very low specificity at the beginning that compensates for a high sensitivity, then the specificity starts to increase again after involving 25 or more proteins.

**Fig 10.**
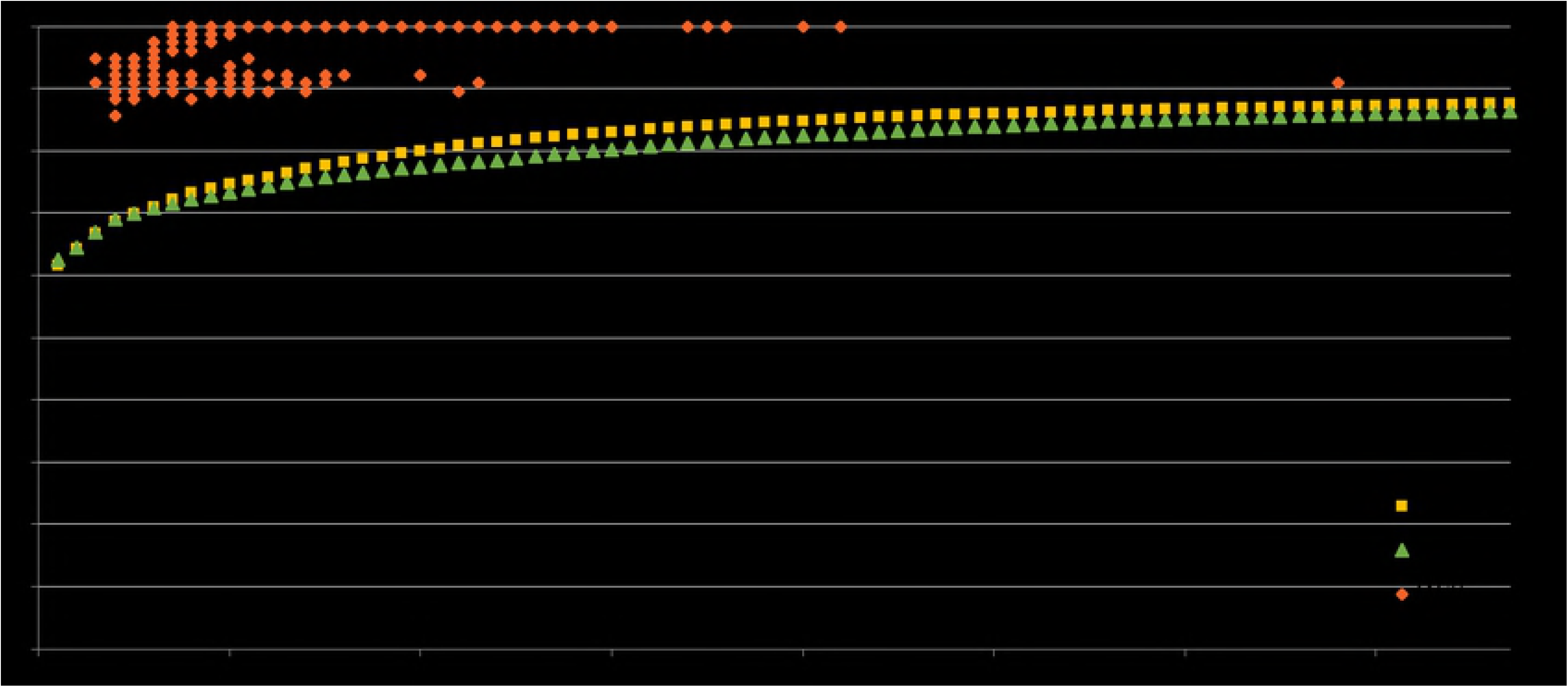
SVM Classification system performance in trisomy mice using different number of features sorted by (a) Fisher, (b) Chi scores.

The Chi test was done on the trisomy mice as shown in appendix 2. As shown, KCG had the higher Chi score, and SOD1 came in the second place, followed by CaNA and ERK. However, this means that SOD1 and CaNA are expressed in both trisomy and control mice. The SVM classification system response using different number of proteins sorted by their Chi score is shown in Fig 10 (b).

Applying the correlation test on the trisomy data showed that only 19 proteins are involved in the subset that obtained a heuristic merit of 0.655 as shown in appendix 2. These proteins are: CAMKII, ERK, PKCAB, BRAF, SOD1, P38, pMTOR, RAPTOR, pP70S6, NUMB, pGSK3B, pPKCG, AcetylH3K9, ARC, IL1B, PSD95, Ubiquitin, BCL2, CaNA. Note that the NUMB, CaNa, pPKCG and SOD1 were also involved in the control mice. This means that trisomy mice show a partial control response to learning. The SVM classification system performance obtained a mean accuracy of 92.3%, a sensitivity of 94.1%, and a specificity of 98.2%.

In trisomy DFS, the same parameters of network configurations of control mice were used. The training data consists of 357, validation data is 77, and a test data of 77 instances. The weights of the additional input layer are illustrated in appendix 2, where 21 proteins have non-zero weights. After applying the 5% of the maximum value threshold, 9 proteins were selected such as: pCAMKII, APP, SOD1, P38, RAPTOR, NUMB, GluR3, GluR4, pGSK3B Tyr216, obtaining an accuracy of 99%, sensitivity of 99% and specificity of 100%. This means that proteins in trisomy are highly distinguishable using DFS and that the effect of memantine injection on the learning is greater in trisomy mice.

The comparison of the proposed techniques for trisomy mice is shown in Fig 11. The DFS showed a very high performance accuracy of 99% at low features while the Fisher and Chi using SVM obtained only 80% and 78% accuracies. This is a proof of nonlinearity in the data classification.

**Fig 11.**
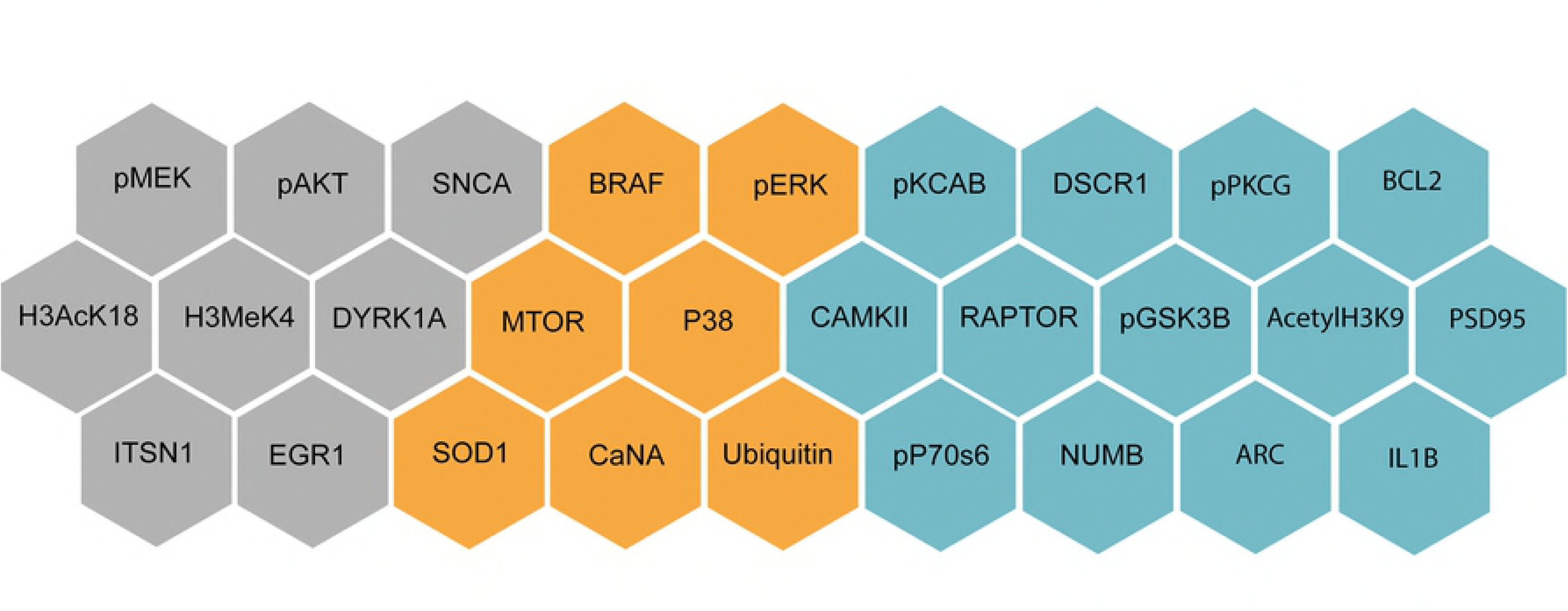
Performance of classification using different selection techniques in trisomy mice.

Similarly, we Compare the proteins selected by the DFS and [5] in trisomy mice, fig 12 shows the proteins selected by DFS, and SOM as well as the common proteins. Among the selected proteins BRAF,pERK, MTOR,P38,SOD1, CaNA, and Ubiquitin were common, whereas pPKCG was selected by our approach which agrees with the results of [35]. To evaluate our selected proteins, linear SVM was used with five cross-validation. Fig 13 shows the results of the five validation iterations. The proteins selected in our study show close variance in the accuracies to [5]’s proteins. The maximum and average accuracies obtained are 91.23% and 88.07% respectively compared to 87.72% and 85.44% from [5]. Therefore, our selected proteins perform a better distinction between the two trisomy mice groups that are with and without learning experience that account for the effect of memantine.

**Fig 12.**
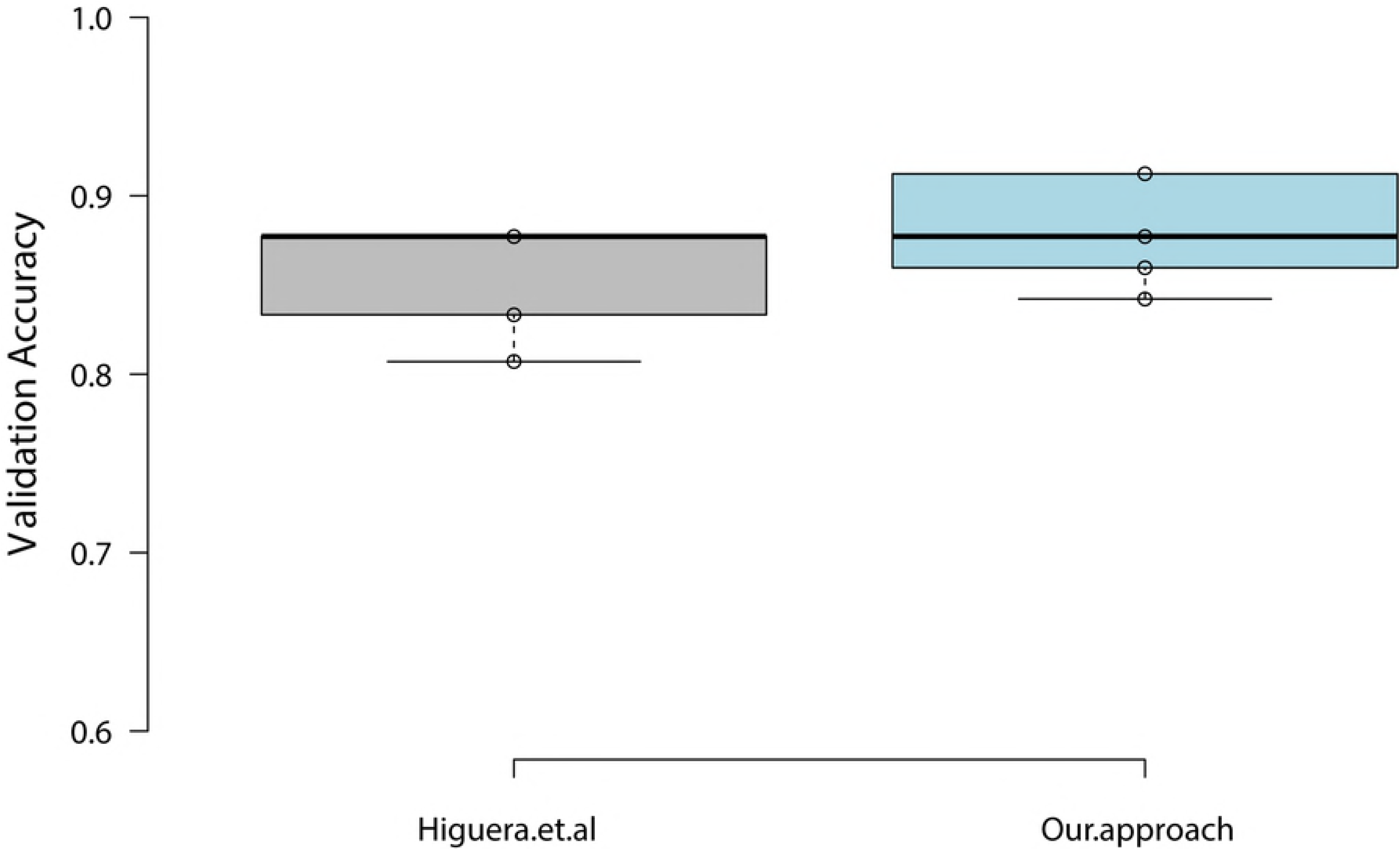
Proteins related to memory and learning from [5] and our approach in trisomy mice group. Gray: proteins selected by [5]. Blue: proteins selected by DFS. Orange: common proteins.

**Fig 13.**
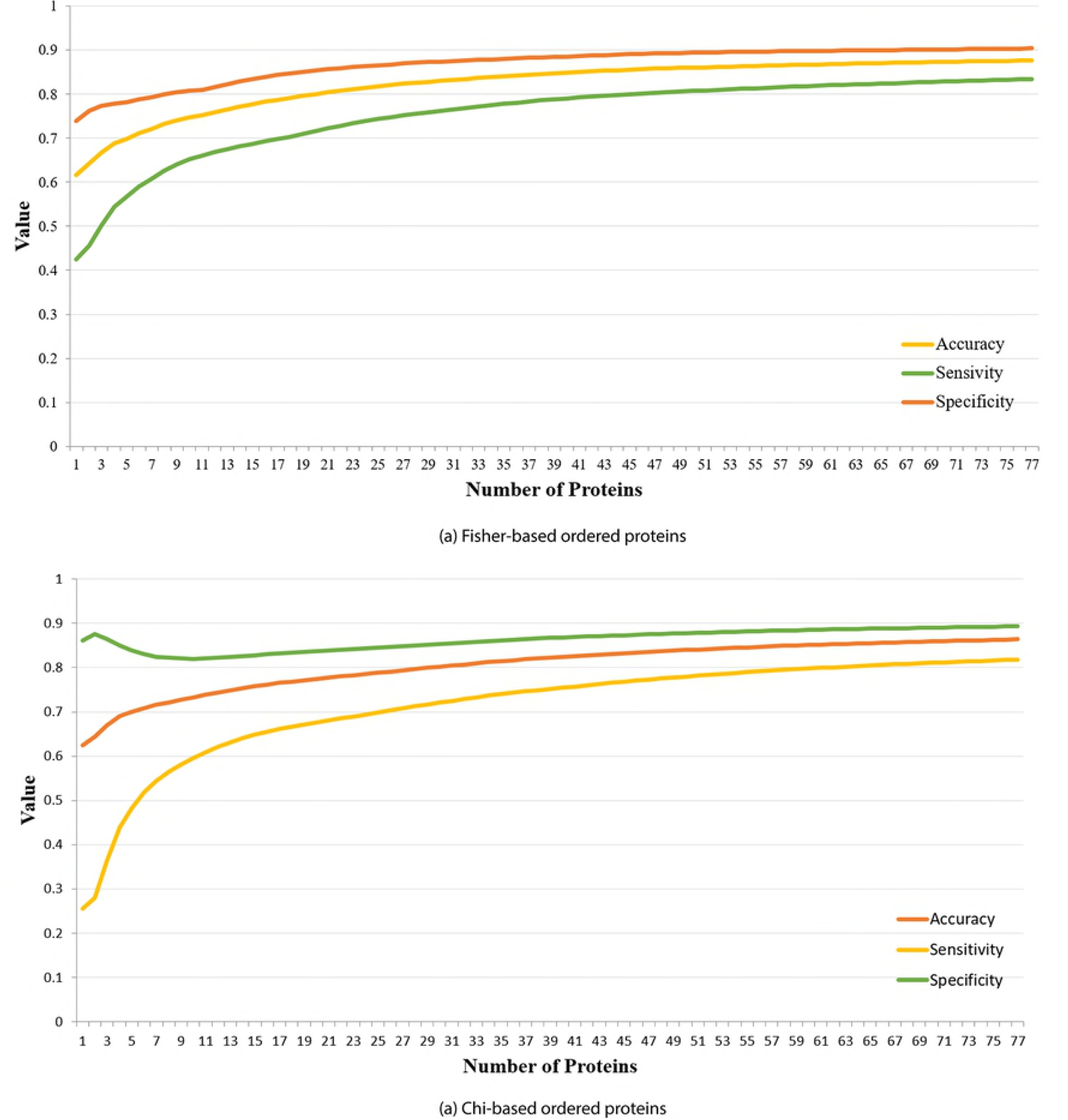
Boxplot of the SVM five cross-validation using the selected proteins in Control mice.

## Control-Trisomy Proteins Analysis

In this experiment, we studied both control and trisomy mice. The dataset was divided into two categories: learned mice and failed to learn mice including the no-learning classes resulted from shock context. The fisher scores of the 77 proteins are shown in appendix 3. The CaNA and SOD1, obtained the highest scores, followed by Ubiquitin and ACR proteins. Not to mention that these proteins had high scores in both control and trisomy mice, which validates the results in the previous sections. Fig 14 (a) shows the SVM system performance versus the number of selected proteins to classify learned and not-learned mice in the whole dataset. The accuracy starts to settle after the first 20 proteins having 85% accuracy, 70% sensitivity, and 84% specificity.

**Fig 14.**
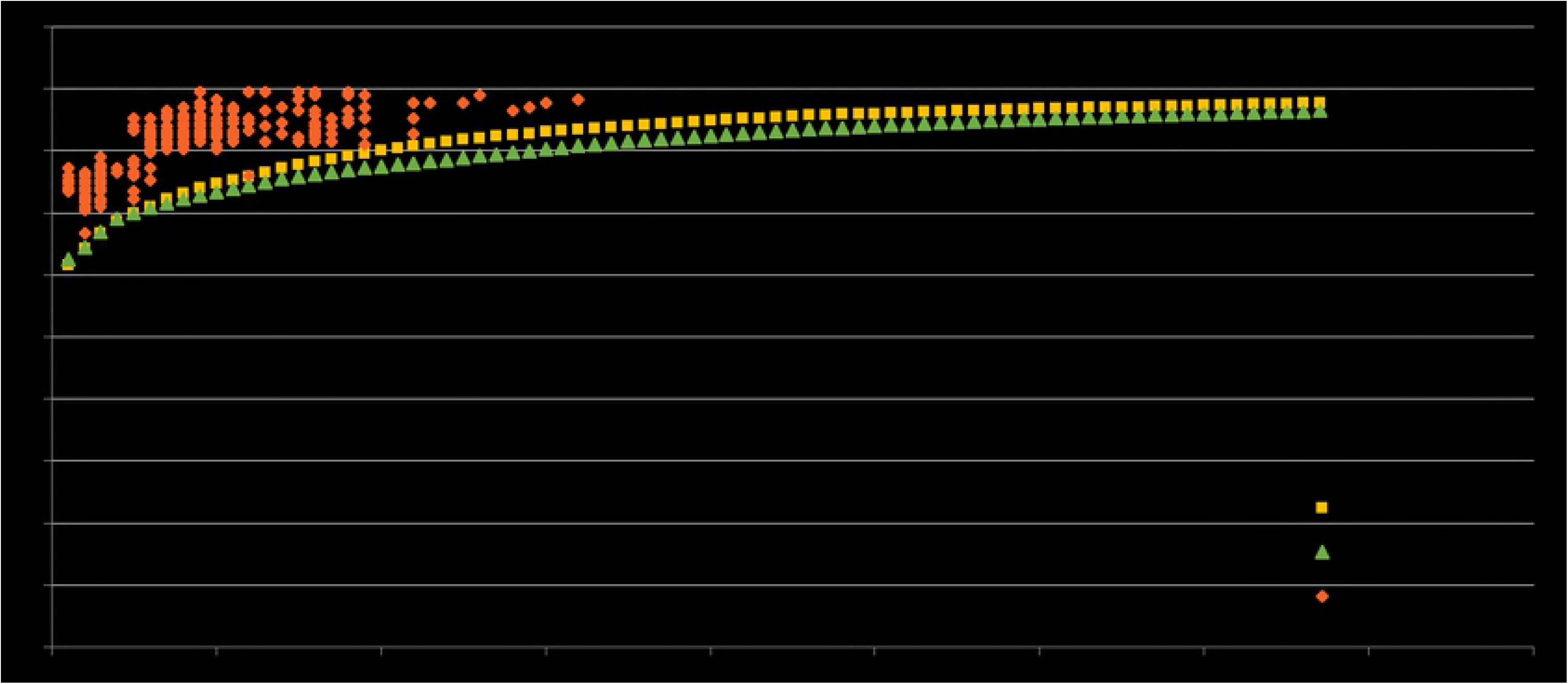
SVM Classification system performance in control-trisomy mice using different number of features sorted by (a) Fisher, (b) Chi scores.

The chi score of each protein is also shown in appendix 3. The CaNA protein held the forth place while SOD1 obtained the fifth place. The highest valued was scored by the H3MeK4 protein. The classification performance in shown in Fig 14 (b). The overall system accuracy is not good, as the maximum accuracy of 86%, a sensitivity of 83% and specificity of 90% were obtained using the full protein set. However, the system accuracy starts to settle after the first 33 first proteins.

The correlation-based protein selection method has obtained a best merit of 0.24 having a set of 20 proteins: pAKT, pCAMKII, pELK, pRSK, P38, DSCR1, NR2B, pNUMB, pP70S6, NUMB, pGSK3B, AcetylH3K9, nNOS, Tau, Ubiquitin, BCL2, H3AcK18, EGR1, H3MeK4, CaNA as shown in appendix 3. Again, CaNA, and NUMB proteins were indicated as significant proteins in the learning process.

In DFS, a train data of 756 instance, 86 validation instances, and 86 test instances were utilized. We used the same parameters of the control and trisomy experiments. The tested model output seven proteins: NR1, pCAMKII, pPKCAB, APP, pNUMB, nNOS, and pGSK3B (Tyr216), where all the scores are shown in appendix 3. The system accuracy is 84% which is considerably low. This might be a result of the non-adjusted network parameters. However, the feature selection, using the specified range of *λ*_1_ from 0 to 0.3 with a step of 0.001, results in an overall high accuracy of more than 85% using seven proteins as shown in fig 15.

**Fig 15.**
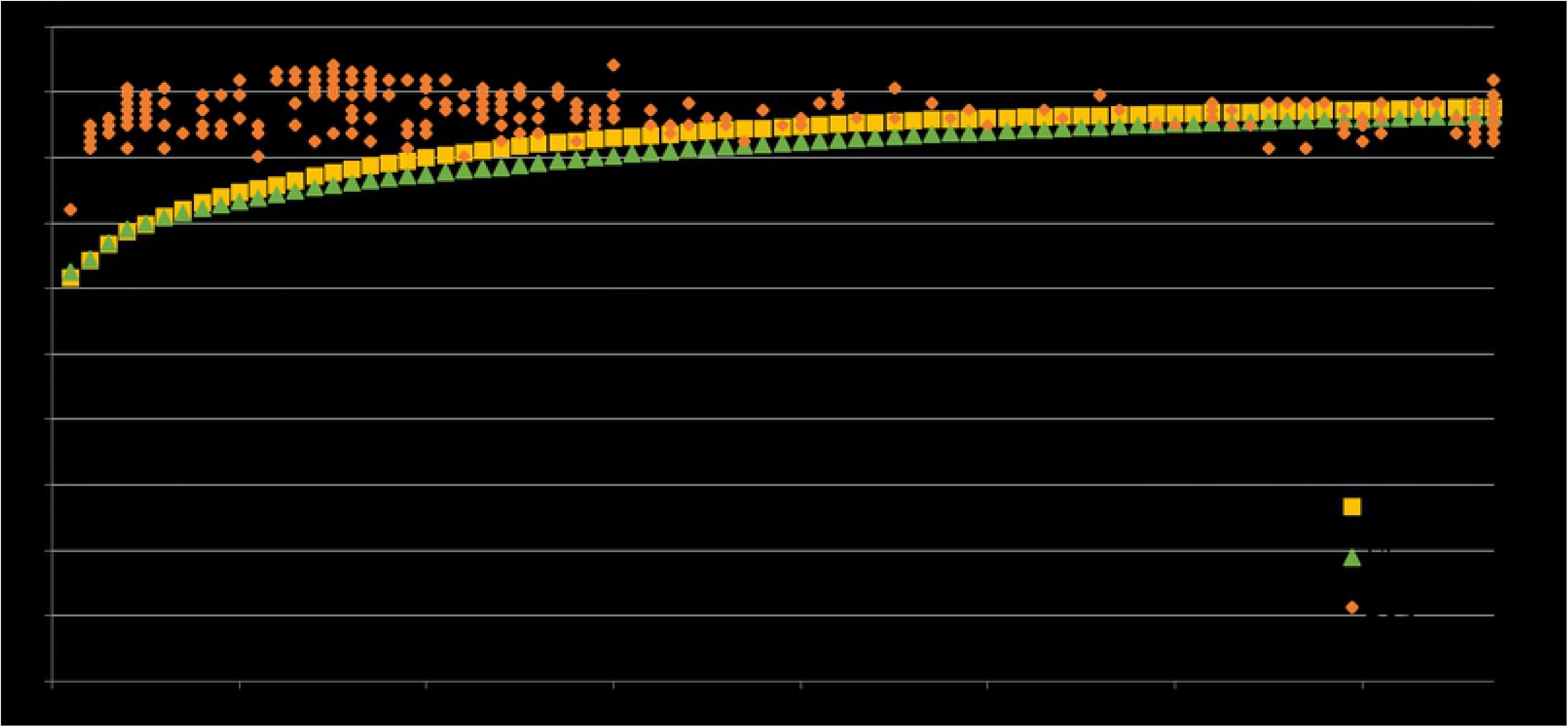
Performance of classification using different selection techniques in control-trisomy mice.

## Conclusion

Down syndrome is a chromosomal condition that is associated with intellectual disability and related to memory and learning deficiency. This disease is complicated and invloves so many genes and pathways. This pathway disturbance can be interpreted from experiments that measure the proteins. In this work, we applied different protein selection techniques in order to select the important proteins that influence the memory and learning among 77 proteins from a Context Fear Shock experiment done on control and trisomy wild mice. We utilized different approaches such as Fisher score, Chi score, correlation-based feature selection using best subset searching algorithm, and deep proteins selection using deep neural networks. In our approach, we compared learning or rescued classes versus no-learning or failed to learn classes. Our study of the proteins in the control mice group indicated that SOD1, CaNA, Ubiquitin, and nNOS proteins have the highest impact on memory and learning process. Utilizing the deep feature selection using two hidden layers, fifteen proteins were selected obtaining an accuracy of 93%. In the trisomy mice case, nine proteins were selected obtaining an accuracy of 99%, indicating that memantine injected to the trisomy mice can promote the response to the learning process. On the other hand, control-trisomy deep feature selection resulted in seven proteins achieving accuracy of 84%. The deep feature selection proved that higher order protein selection is needed in the prediction of significant proteins related to memory and learning, where it outperformed the Fisher, Chi, and correlation-based protein selection methods. Moreover, the proteins selected in this research showed a higher distinction ability between each of the two groups involved in the classification. In addition, the deep feature selection showed a better performance compared to SOM, since it take advantage of using the data label making it a supervised learning.

Freepik Graphics;. https://www.freepik.com/free-vectors/templates.

## References

1. Weijerman ME, de Winter JP. Clinical practice The care of children with Down syndrome. Consequences of Down syndrome for patient and family. 2011;169:11.

2. World Health Organization GRC. WHO Genes and human disease; 2010.

3. Irving C, Basu A, Richmond S, Burn J, Wren C. Twenty-year trends in prevalence and survival of Down syndrome. European Journal of Human Genetics. 2008;16(11):1336.

4. Sturgeon X, Gardiner KJ. Transcript catalogs of human chromosome 21 and orthologous chimpanzee and mouse regions. Mammalian Genome. 2011;22(5-6):261–271.

5. Higuera C, Gardiner KJ, Cios KJ. Self-organizing feature maps identify proteins critical to learning in a mouse model of down syndrome. PloS one. 2015;10(6):e0129126.

6. Sturgeon X, Le T, Ahmed MM, Gardiner KJ. Pathways to cognitive deficits in Down syndrome. In: Progress in brain research. vol. 197. Elsevier; 2012. p. 73–100.

7. Reeves RH, Irving NG, Moran TH, Wohn A, Kitt C, Sisodia SS, et al A mouse model for Down syndrome exhibits learning and behaviour deficits. Nature genetics. 1995;11(2):177.

8. Allore R, O’Hanlon D, Price R, Neilson K, Willard H, Cox D, et al Gene encoding the beta subunit of S100 protein is on chromosome 21: implications for Down syndrome. Science. 1988;239(4845):1311–1313.

9. Siddiqui A, Lacroix T, Stasko MR, Scott-McKean JJ, Costa AC, Gardiner KJ. Molecular responses of the Ts65Dn and Ts1Cje mouse models of Down syndrome to MK-801. Genes, Brain and Behavior. 2008;7(7):810–820.

10. Shim KS, Lubec G. Drebrin, a dendritic spine protein, is manifold decreased in brains of patients with Alzheimer’s disease and Down syndrome. Neuroscience letters. 2002;324(3):209–212.

11. Block A, Ahmed MM, Dhanasekaran AR, Tong S, Gardiner KJ. Sex differences in protein expression in the mouse brain and their perturbations in a model of Down syndrome. Biology of sex differences. 2015;6(1):24.

12. Stagni F, Giacomini A, Guidi S, Ciani E, Bartesaghi R. Timing of therapies for Down syndrome: the sooner, the better. Frontiers in behavioral neuroscience. 2015;9:265.

13. Furqan MS, Siyal MY. Protein Map of Control Mice Exposed to Context Fear Using A Novel Implementation of Granger Causality. In: Artificial Intelligence, Modelling and Simulation (AIMS), 2015 3rd International Conference on. IEEE; 2015. p. 96–98.

14. Ahmed MM, Dhanasekaran AR, Block A, Tong S, Costa AC, Stasko M, et al Protein dynamics associated with failed and rescued learning in the Ts65Dn mouse model of Down syndrome. PloS one. 2015;10(3):e0119491.

15. Boada R, Hutaff-Lee C, Schrader A, Weitzenkamp D, Benke T, Goldson E, et al Antagonism of NMDA receptors as a potential treatment for Down syndrome: a pilot randomized controlled trial. Translational psychiatry. 2012;2(7):e141.

16. Costa AC, Scott-McKean JJ, Stasko MR. Acute injections of the NMDA receptor antagonist memantine rescue performance deficits of the Ts65Dn mouse model of Down syndrome on a fear conditioning test. Neuropsychopharmacology. 2008;33(7):1624.

17. Kohavi R, John GH. Wrappers for feature subset selection. Artificial intelligence. 1997;97(1-2):273–324.

18. John GH, Kohavi R, Pfleger K. Irrelevant features and the subset selection problem. In: Machine Learning Proceedings 1994. Elsevier; 1994. p. 121–129.

19. Skalak DB. Prototype and feature selection by sampling and random mutation hill climbing algorithms. In: Machine Learning Proceedings 1994. Elsevier; 1994. p. 293–301.

20. Mitchell M, Holland JH, Forrest S. When will a genetic algorithm outperform hill climbing. In: Advances in neural information processing systems; 1994. p. 51–58.

21. Austin S, Schwartz R, Placeway P. The forward-backward search algorithm. In: Acoustics, Speech, and Signal Processing, 1991. ICASSP-91., 1991 International Conference on. IEEE; 1991. p. 697–700.

22. Brownlee J. Feature selection to improve accuracy and decrease training time. Machine Learning Mastery. 2014;.

23. Yu L, Liu H. Feature selection for high-dimensional data: A fast correlation-based filter solution. In: Proceedings of the 20th international conference on machine learning (ICML-03); 2003. p. 856–863.

24. Hall MA. Correlation-based feature selection of discrete and numeric class machine learning. 2000;.

25. Li Y, Chen CY, Wasserman WW. Deep feature selection: theory and application to identify enhancers and promoters. Journal of Computational Biology. 2016;23(5):322–336.

26. Longford NT. A fast scoring algorithm for maximum likelihood estimation in unbalanced mixed models with nested random effects. Biometrika. 1987;74(4):817–827.

27. Univesity of Pennsylvania LD. utorial: Pearson’s Chi-square Test for Independence; 2016.

28. Hall MA. Correlation-based feature selection for machine learning. 1999;.

29. Hornik K, Stinchcombe M, White H. Multilayer feedforward networks are universal approximators. Neural networks. 1989;2(5):359–366.

30. Fletcher R, Powell MJ. A rapidly convergent descent method for minimization. The computer journal. 1963;6(2):163–168.

31. Cortes C, Vapnik V. Support-vector networks. Machine learning. 1995;20(3):273–297.

32. Aizerman MA. Theoretical foundations of the potential function method in pattern recognition learning. Automation and remote control. 1964;25:821–837.

33. Murakami K, Murata N, Noda Y, Tahara S, Kaneko T, Kinoshita N, et al SOD1 (copper/zinc superoxide dismutase) deficiency drives amyloid β protein oligomerization and memory loss in mouse model of Alzheimer disease. Journal of Biological Chemistry. 2011;286(52):44557–44568.

34. Havekes R, Nijholt IM, Luiten PG, Van der Zee EA. Differential involvement of hippocampal calcineurin during learning and reversal learning in a Y-maze task. Learning & Memory. 2006;13(6):753–759.

35. Giese KP, Mizuno K. The roles of protein kinases in learning and memory. Learning & memory. 2013;20(10):540–552.

36. Freepik Graphics;. https://www.freepik.com/free-vectors/templates.

